# Automating the analysis of fish abundance using object detection: optimising animal ecology with deep learning

**DOI:** 10.1101/805796

**Authors:** Ellen M. Ditria, Sebastian Lopez-Marcano, Michael K. Sievers, Eric L. Jinks, Christopher J. Brown, Rod M. Connolly

## Abstract

Aquatic ecologists routinely count animals to provide critical information for conservation and management. Increased accessibility to underwater recording equipment such as cameras and unmanned underwater devices have allowed footage to be captured efficiently and safely. It has, however, led to immense volumes of data being collected that require manual processing, and thus significant time, labour and money. The use of deep learning to automate image processing has substantial benefits, but has rarely been adopted within the field of aquatic ecology. To test its efficacy and utility, we compared the accuracy and speed of deep learning techniques against human counterparts for quantifying fish abundance in underwater images and video footage. We collected footage of fish assemblages in seagrass meadows in Queensland, Australia. We produced three models using a MaskR-CNN object detection framework to detect the target species, an ecologically important fish, luderick (*Girella tricuspidata*). Our models were trained on three randomised 80:20 ratios of training:validation data-sets from a total of 6,080 annotations. The computer accurately determined abundance from videos with high performance using unseen footage from the same estuary as the training data (F1 = 92.4%, mAP50 = 92.5%), and from novel footage collected from a different estuary (F1 = 92.3%, mAP50 = 93.4%). The computer’s performance in determining MaxN was 7.1% better than human marine experts, and 13.4% better than citizen scientists in single image test data-sets, and 1.5% and 7.8% higher in video data-sets, respectively. We show that deep learning is a more accurate tool than humans at determining abundance, and that results are consistent and transferable across survey locations. Deep learning methods provide a faster, cheaper and more accurate alternative to manual data analysis methods currently used to monitor and assess animal abundance. Deep learning techniques have much to offer the field of aquatic ecology.

## 1. Introduction

The foundation for all key questions in animal ecology revolves around the abundance, distribution and behaviour of animals. Collecting robust, accurate and unbiased information is therefore vital to understanding ecological theories and applications. Many of the invasive data collection methods traditionally used to collect this information in animal ecology, such as tagging, netting and trawling, are now largely unnecessary due to remote data collection using cameras. The development and availability of these devices have facilitated more accurate and cheaper methods of data collection, with reduced risk to the operator (Hodgson et al. 2013). Most importantly from a scientific perspective, they have increased sampling accuracy as well as replicability and reproducibility (Weinstein 2017), which form the basis of a sound scientific study (Leek & Peng 2015). However, the amount of data now being generated can be overwhelming. The solution has become the new problem.

Much like the physical collection of data, manual processing of data is often labour-intensive, time-consuming and extremely costly (Weinstein 2017). This has led to invaluable data collected over large temporal and spatial scales laying unused in storage libraries. In Australia, for example, the Integrated Marine Observing System (IMOS) collects millions of images of coral reefs every year, yet despite affiliations and partnerships with a range of universities and management agencies, less than 5% of these are analysed by experts (Moniruzzaman et al. 2017). This apparently never-ending stream of data brings a new challenge for ecologists; to find or develop the analytical tools needed to extract information from the immense volumes of incoming images and video content (Valletta et al. 2017).

Fortunately, recent advances in machine learning technologies have provided one such tool to help combat this problem; deep learning. Deep learning is a subset of machine learning consisting of a number of computational layers within an architectural framework designed to process data that is difficult to model analytically, such as raw images and video footage (LeCun et al. 2015). Although neural networks are not a new technology (Rawat & Wang 2017), the relatively recent advances in graphics processing units (GPUs) have spurred an increase in their application for computer vision data. In the CNN, data are fed into an input layer, while an output layer is sorted into categories pre-determined by manual training, in a process known as supervised learning (Rawat & Wang 2017).

Although deep learning techniques are being implemented enthusiastically in terrestrial ecology, it is currently an under-exploited tool in aquatic environments (Moniruzzaman et al. 2017, Xu et al. 2019). As the global challenges in marine science and management increase (Halpern et al. 2015), it is critical for marine science to realise the potential automated analysis offers (Malde et al. 2019). Relative to terrestrial environments, however, obtaining useable footage in marine environments to achieve acceptable computational performance presents a unique set of challenges. For example, there are often high levels of environmental complexities in marine environments which can interfere with clear footage, including variable water clarity, complex background structures, decreased light at depth, and obstruction due to schooling fish (Mandal et al. 2018, Salman et al. 2019). Although these factors may affect the quality of images and videos, deep learning methods have proven successful in a range of marine applications (Galloway et al. 2017, Arellano-Verdejo et al. 2019).

Efforts to use deep learning methods in marine environments currently revolve around the automated *classification* of specific species. Attempts to classify tropical reef fish have achieved high levels of performance and have also outperformed humans in species recognition (Villon et al. 2018). There have also been suggestions from classification studies on freshwater fish to incorporate other strategies for increasing performance, such as including taxonomic family and order (dos Santos & Gonçalves 2019). Although all marine environments have challenging conditions, the tropical reef studies by Villon et al. (2018) and Salman et al. (2019) typically operate with high visibility, high fish abundance, and highly variable inter-specific morphology, which makes distinguishing different species easier (Xu & Matzner 2018). Conversely, coastal and estuarine systems often suffer poor visibility due to complex topography, anthropogenic eutrophication, and sediment induced turbidity (Lehtiniemi et al. 2005, Baker & Sheaves 2006, Lowe et al. 2015).

Although classification enables the determination of species, its usefulness for answering broad ecological questions is rather limited. Object detection allows us to classify both *what* is in a frame and *where* it is and therefore enables us to determine both the species in an area and their abundance (eg. Maire et al. 2015, Salberg 2015, Gray et al. 2019b).

Here, we use fish inhabiting subtropical seagrass meadows as a case study to explore the viability of computer vision and deep learning as a suitable, non-invasive technique using remotely collected data in a variable marine environment. Seagrass meadows provide critical ecosystem services such as carbon sequestration, nutrient cycling, shoreline stabilisation and enhanced biodiversity (Waycott et al. 2009, Sievers et al. 2019). However, many seagrass meadows are being lost and degraded due to a range of anthropogenic stressors, such as overfishing, eutrophication and physical disturbances (Orth et al. 2006). Due to their background complexity, constant movement, and ability to obscure fish, seagrass may prove to be a difficult habitat to implement a deep learning solution. Luderick (*Girella tricuspidata*) is a common herbivorous fish found along the east coast of Australia and is abundant in coastal and estuarine systems, including seagrass meadows (Ferguson et al. 2013). Unlike most herbivorous fish in seagrass meadows, this species grazes on both the epiphytic algae that grows on seagrass and the seagrass itself, making it of interest ecologically (Gollan & Wright 2006). Using this ecologically important ecosystem, we specifically aim to deduce whether deep learning techniques can be used to determine: (1) the accurate object detection of a target species, (2) the flexibility of algorithms in analysing data across locations, and (3) the comparative performance between computers and humans in determining abundance from images and video footage. As far as we are aware, this is the first time that humans and deep learning algorithms have been compared in their ability to quantify abundance from underwater video footage, or that object detection and computer vision methods have been used in estuarine systems.

## 2. Methods

### 2.1 Training data-set

We used submerged action cameras (Haldex Sports Action Cam HD 1080p) to collect video footage of luderick in the Tweed River estuary in southeast Queensland (−28.169438, 153.547594), between February and July 2019. Each sampling day, six cameras were deployed for 1 h over a variety of seagrass patches; the angle and placement of cameras was varied among deployment to ensure a variety of backgrounds and fish angles. Videos were trimmed for training to contain only footage of luderick and split into 5 frames per second.

### 2.2 Convolutional Neural Network

The object detection framework we used is an implementation of Mask R-CNN developed by Massa & Girshick (2018). Mask R-CNN works by classifying and localising the region of interest (RoI). It extends previous frameworks in that it can predict a segmentation mask on the RoI, and currently has the highest performance output for deep learning models (He et al. 2017, Dai et al. 2019). To develop our model, we used a ResNet50 configuration, pre-trained on the ImageNet-1k data-set. This configuration provides an acceptable balance between training time and performance (Massa & Girshick 2018). We conducted the model training, testing and prediction tasks on a Microsoft Azure Data Science Virtual Machine powered by an NVIDIA V100 GPU. Data preparation and annotation tasks were carried out using software developed at Griffith University. While deep learning has begun to be adopted for ecological data analysis in the last two years, its use in the environmental sciences requires substantial software engineering knowledge, as unfortunately there is not yet an accessible software package for ecologists (Piechaud et al. 2019). The development of this interface for manual annotation, that can be retrained for different species, takes strides towards an end-to-end, user-friendly application tailored for ecologists. A trained team in fish identification manually drew segmentation masks around luderick (i.e. our RoI, Fig. 2.1) and annotated 6,080 fish for the training data-set. Luderick were annotated if they could be positively identified at any time within the video the image came from.

**Fig. 1.**
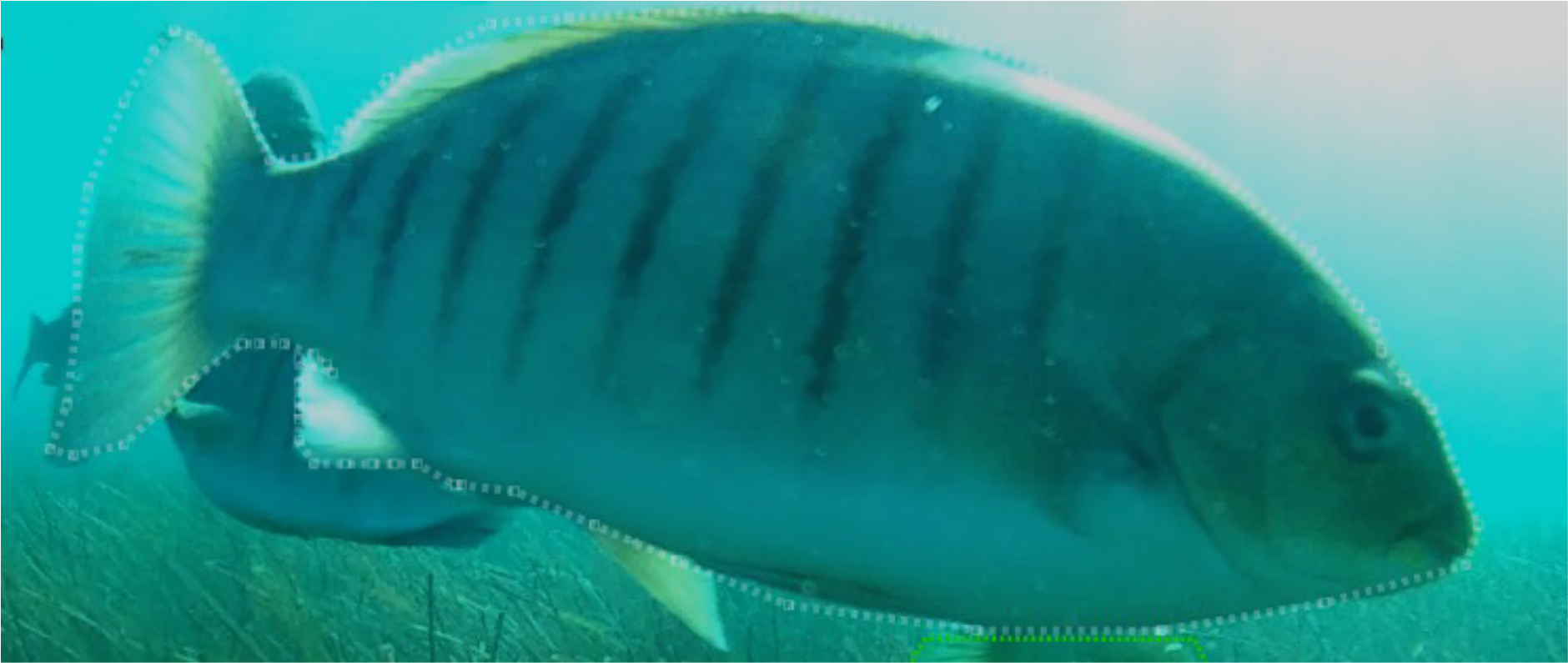
Training data-set image demonstrating manual segmentation mask (white dashed line around fish) denoting the region of interest (RoI).

**Fig. 2.**
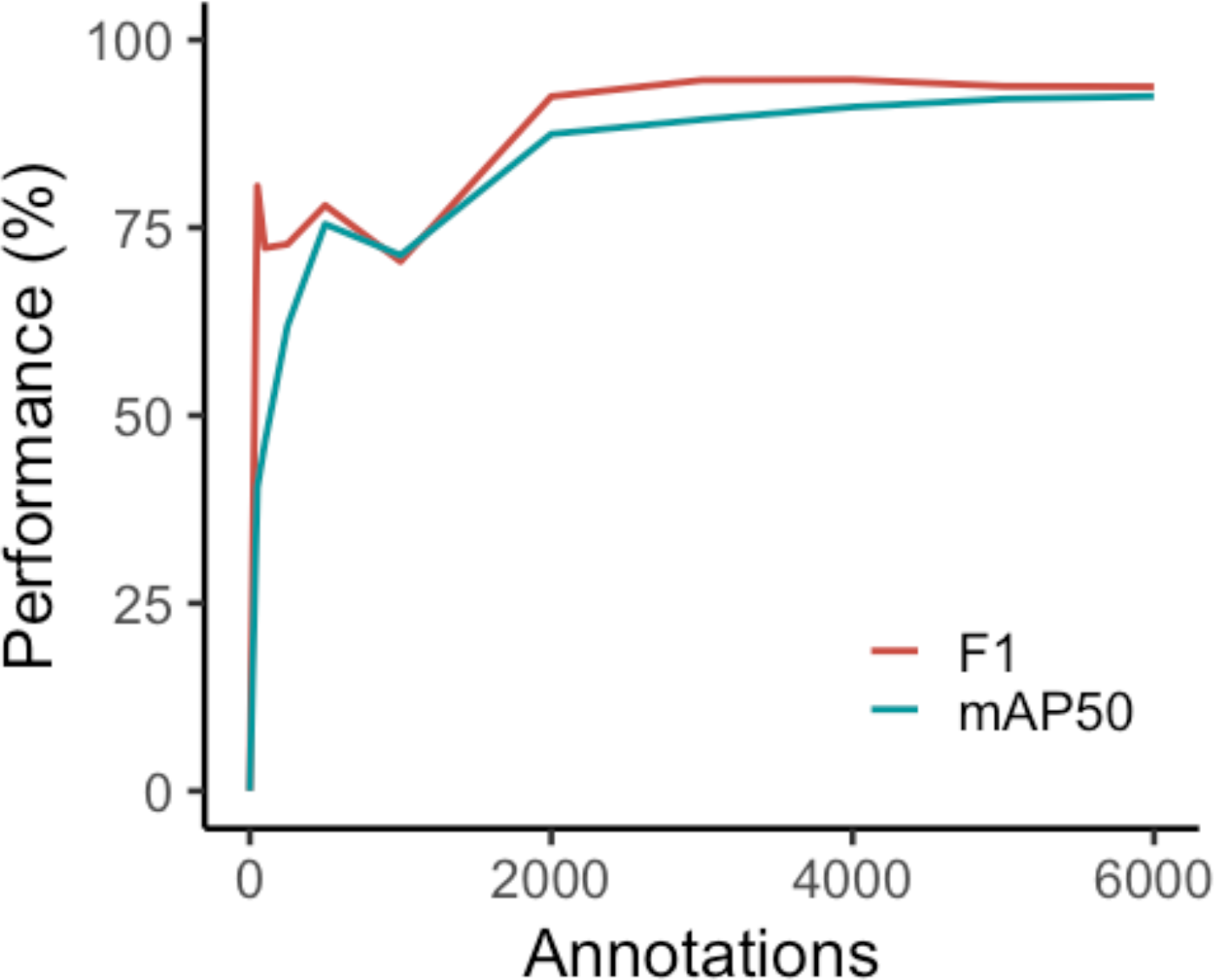
Performance curve showing the computer’s ability to fit a segmentation mask around the luderick (performance scored by mAP50) and in identifying abundance (performance scored by F1).

The utility of the model depends on how accurately the computer identifies the presence of luderick, which we quantified in two ways based on the interactions between precision (P) and recall (R). Precision is how rigorous the model is at identifying the presence of luderick, and recall is the number of the total positives the model captured (Everingham et al. 2010). Generally, an increase in recall results in decreased precision and vice versa and were calculated as follows:

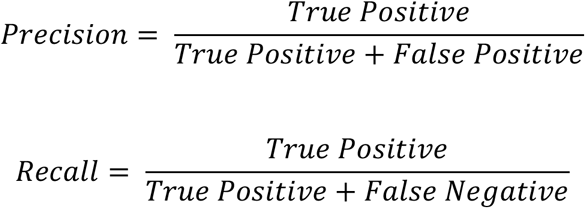

Firstly, the computer’s ability to fit a segmentation mask around the RoI was determined by the mean average precision value (mAP) (Everingham et al. 2010).

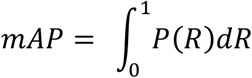

We used the mAP50 value in this study, which equates to how well the model overlapped a segmentation mask around at least 50% of the ground truth outline of the fish. The higher this value, the more accurate the model was at overlapping the segmentation mask. Secondly, the success of our model in answering ecological questions on abundance was determined by an F1 score:

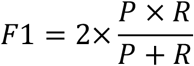

We used the F1 score and mAP50 values to assess the performance of the computer model. All predictions were made with a confidence threshold of 90%, that is, the algorithm was at least 90% sure that it was identifying a luderick to minimise the occurrence of false negatives. This threshold was chosen as it typically maximised F1 performance by filtering out false positives.

### 2.3 Model Validation and Performance Curve

Models were trained using a random 80% sample of the annotated dataset, with the remaining 20% used to form a validation dataset (Alexandropoulos et al. 2019). Training performance was then measured against the validation set to monitor for overfitting. Overfitting is a phenomenon when the computer becomes dependent on, and memorises the training data, failing to perform well when tested on data it has not encountered previously (Chicco 2017). We minimised overfitting by using the early-stopping technique (Prechelt 1998). In our case, this was achieved by assessing the mAP50 on the validation set at intervals of 2,500 iterations and determined where the performance began to drop (Chicco 2017). The same computer algorithm was used to train three different models on three different randomised 80/20 subsets of the whole training data set to account for variation in the training and validation split. These models were subsequently used to compare the unseen and novel test data-set, and in the human vs computer test.

We generated a performance curve to confirm that variation among models was sufficiently low to ensure consistency in in performance across the three models. Random subsets of still images were selected from the training data-set. These subsets of data increased in volume to determine the performance of the model as training data increase. As the volume of training data increased, the risk of overfitting decreased so the number of training iterations were adjusted to maintain optimum performance.

Manual annotation cost can be a significant factor to consider when training CNN networks and can also be monitored by using the performance curve. Time stamps were added to the training software to record the speed at which training data was annotated to infer total annotation time of the training data by humans. We used this data to determined how much training is required by this model to produce high accuracy, and thus also the effort needed to produce a consistent and reliable ecological tool.

### 2.4 Model performance

The 80/20 validation test is an established method in machine learning to assess the expected performance of the final model (Alexandropoulos et al. 2019). However, using deep learning to answer ecological questions requires another testing procedure to accurately reflect the usability of the model when analysing new data. We therefore also tested the model against annotations from two types of new footage not used for the training data-set. We used unseen footage from the same location in the Tweed River estuary (‘Unseen’), as well as from a novel location (‘Novel’), being seagrass meadows in a separate estuary system in Tallebudgera Creek (−28.109721, 153.448975). A t-test was used to compare the performance of the three models between the unseen test-set from Tweed estuary, and the novel test-set from Tallebudgera.

### 2.5 Human vs Computer

Creating an automated data analysis system aims to lessen the manual workload of humans by creating a faster, yet accurate, alternative. Therefore, it is crucial to not only know how well the model performs, but also to assess its capabilities in speed and accuracy, compared to current human methods. This “human vs computer” method analysis compared Citizen Scientists and Experts against the computer: 1) Citizen Scientists were undergraduate marine science students and interested members of the public (n = 20) 2) Experts were fish scientists with a PhD or currently studying for one (n = 7),, and 3) the computer models (n = 3). We compared these groups using both video footage (n=31) and images (n=50), and analysed differences in test speed and performance. Both the image set and videos were run through the three deep learning models to account for variation in performance in the 80% of training data used to train the models. The number of false negatives, false positives, proportion of accurate answers (observed answers divided by ground truth) as well as the overall F1 score were recorded. Citizen Scientist and Experts were provided with a package that contained a link to the video test uploaded to YouTube, the image set sent as a zip file, instruction sheet, example images of the target species and datasheets. This process was set up to minimise bias in training the human subjects that may have occurred if the test was explained verbally. Humans were instructed to only record the target species if they could visually identify the luderick with confidence. Participants were required to estimate the maximum number of luderick in any single frame per video and per still image (MaxN), simulating the most popular manual method currently used in analysing videos (e.g. Gilby et al. 2017). Start and end time of each test was also recorded to compare how quickly the participants completed the task, compared to the deep learning algorithm. The still image data-set was randomly selected from the “unseen” test video footage and used as the ground truth for images. The video footage was expertly annotated at five frames per second and used as the ground truth for videos. Luderick were only annotated if they could be positively identified at least at one instance in the video. This enabled us to quantitatively compare the human and computer accuracy in determining MaxN, assessed using the overall F1 score for each test.

## 3. Results

### 3.1 Performance curve

Based on the computer algorithm curve, F1 performance began to plateau earlier than mAP50 (Fig. 2.). F1 varied only 0.9% from 2,000 annotations to 6,000 annotations compared to an increase of 3.1% by mAP50 at the same annotations. At lower volumes of training annotations (between 0 and 1,000), the performance of both mAP50 and F1 fluctuated. Even with our streamlined process for annotation, the average time for an operator to annotate one fish was 36 seconds, and the total time to annotate all 6,080 images was in the order of 60 hours.

#### Model performance

Performance was high for both the Unseen and Novel test sets (mAP and F1 both >92%). Based on F1 scores, the computer performed equally well (t-test; t = −0.01, p = 0.99) on the Unseen (92.4%) and novel (92.3%; Fig 3). Similarly, the difference in performance for mAP50 was non-significant (t = 1.4, p = 0.29) on the Unseen (92.5%) and Novel (93.4%) test-sets.

**Fig. 3.**
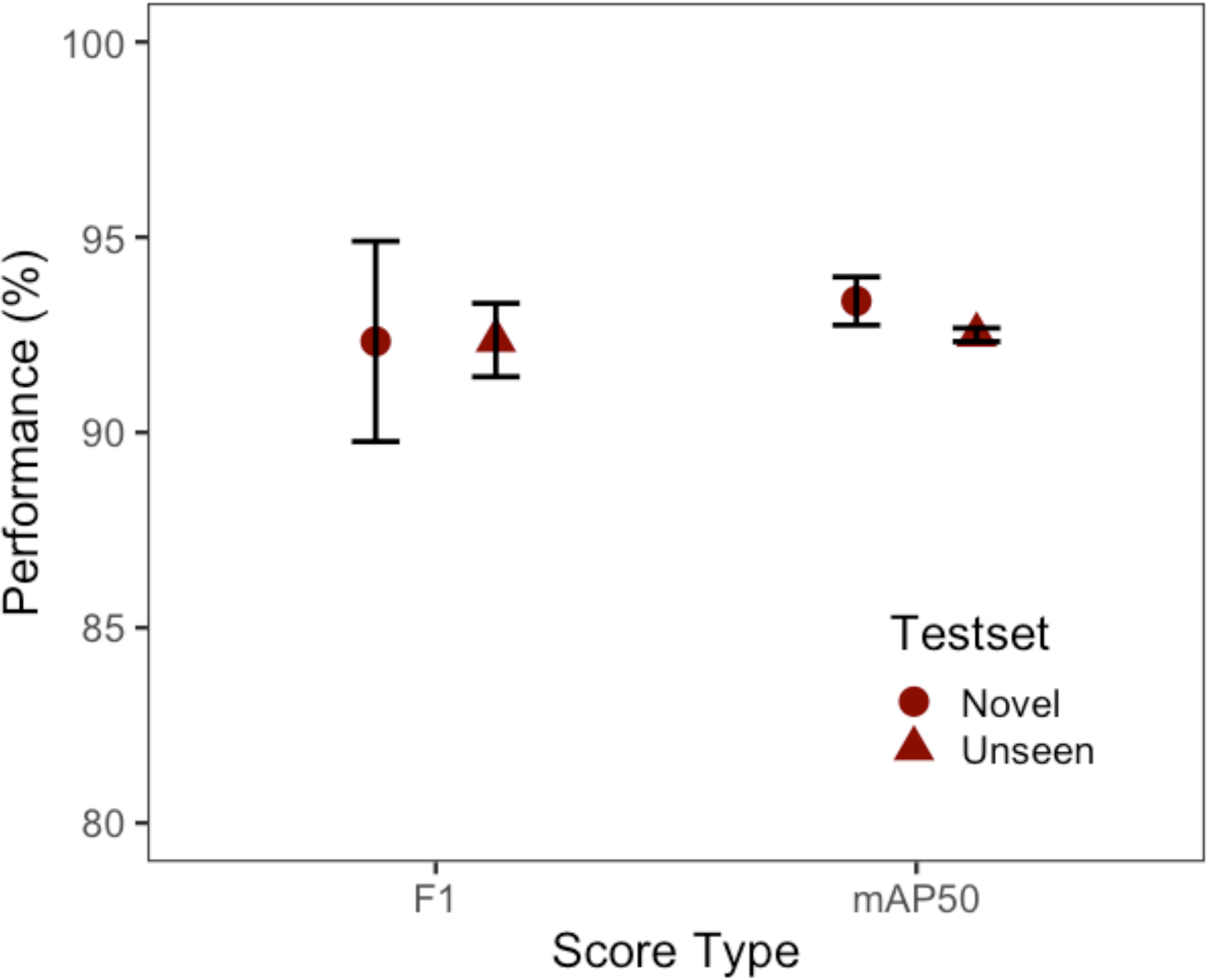
The performance of the three model’s F1 and mAP50 scores (mean, SE) for the unseen test footage from the same location and novel footage (Unseen; 32 videos, Novel; 32 videos).

#### Human vs Machine

The computer algorithm achieved the highest mean F1 score in both the image (95.4%) and the video-based tests (86.8%), when compared with the experts and citizen scientists. The computer also had fewer false positives (incorrectly identifying another species as luderick) and false negatives (incorrectly ignoring a luderick) in the image test. The computer models also had the lowest rate of false positives in the video-based test when compared to both human groups, but had the highest rate of false negatives. The computer performed the task far faster than both human groups. Experts on average performed better (F1) than the citizen scientists in both tests, and had higher accuracy scores (Table 1).

**Table 1.** Summary of performance measures comparing averaged scores from computer vs humans (citizen scientists and experts). Accuracy is displayed as the observer answer divided by the ground truth. Speed is measured as seconds per image, and minutes per minute of video. Images N = 50, Videos N = 31.

| **Analysis Method** | **False Negatives** | **False Positives** | **Accuracy (prop. +/-)** | **F1 (%) (SE)** | **Speed (mins) (SE)** |
| --- | --- | --- | --- | --- | --- |
| **Images** |  |  |  |  |  |
| Citizen Scientist | 28.6 | 7.2 | -0.14 | 82.0 (2.8) | 12.6 (1.4) |
| Expert | 18.1 | 5.6 | -0.08 | 88.3 (8.4) | 14.3 (4.0) |
| Computer | 11.7 | 4.7 | -0.12 | 95.4 (0.9) | 0.4 (0.0) |
| **Videos** |  |  |  |  |  |
| Citizen Scientist | 20.9 | 12.6 | -0.10 | 79.0 (2.4) | 2.4 (2.4) |
| Expert | 12.1 | 11.9 | +0.06 | 85.3 (6.9) | 2.8 (4.4) |
| Computer | 24.3 | 2.7 | -0.10 | 86.8 (1.6) | 1.2 (0.3) |

F1 scores were most variable for the citizen scientist group, with the difference between the lowest and the highest score for the image and video tests being 40.1% and 35.1%, respectively. The computer achieved the lowest variance, with these values only 3.1% for the video test and 1.7% for the image test (Fig. 4).

**Fig. 4.**
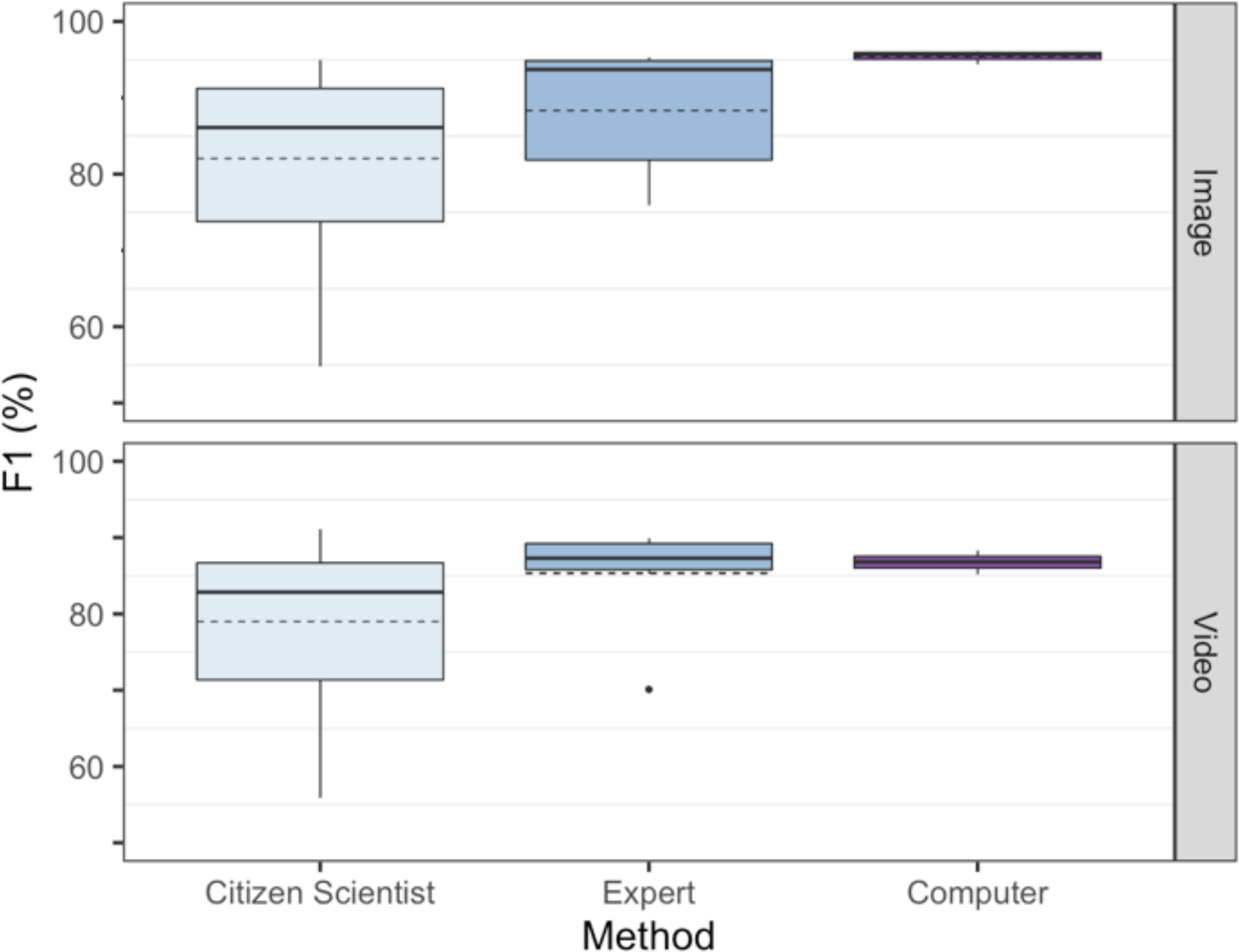
Overall test performance in determining abundance (F1) by Computer vs Humans (Citizen Scientists and Experts) based on identical tests using 50 images and 31 videos. Variance was highest and performance lowest in the citizen scientist group while the computer had the lowest variance and highest performance. Solid line denotes median, dashed line the mean.

## 5. Discussion

Our object detection models achieved high performance on a previously unseen data set, and maintained this performance on footage collected in a novel location. It outperformed both classes of humans (citizen scientists and experts) in speed and performance, with high consistency (i.e. low variability).

We clearly show that our model is fully capable of accurately performing the same on novel footage from locations beyond the data used for training. Few previous demonstrations of the utility of deep learning have tested algorithms under these novel conditions, but is one which consider important for determining how transferable the model is to practising environmental scientists. For our example, our intention was to test how robust and flexible the algorithm was in identifying luderick under different environmental conditions which can vary with tides, water clarity, ambient light, differences in non-target fish species and backgrounds. In a study conducted by Xia et al. (2018) on sea cucumbers, a novel test data set comprised of internet images demonstrated an accuracy of 76.3%. This performance was significantly lower than the test data set the model was trained on which achieved an accuracy of 97.6%. Similarly, Xu and Matzner (2018) attempted to monitor the effects of water turbines on local fish species at three different sites, but their model only generated a 53.9% accuracy. All three sites exhibited their own unique challenges to underwater data collection, including occlusion due to bubbles from fast-flowing water and debris, that made fish detection difficult even for a human observer. Their study demonstrates the aforementioned environmental challenges marine scientist face in using computer vision. Despite the performance limitations of deep learning when provided with limited training data, one reason that our models produced high-performance results from the novel location is the broad variation in environmental conditions and camera angles in the training data. Future work on this topic could extend the novel test to include an even wider array of novel locations to further assess the robustness of the model.

The computer’s high performance, speed and low variance compared to humans suggests that it is a suitable model to replace manual efforts to determine MaxN in marine environments. Deep learning may be the solution for researchers to avoid analytical bottlenecks (Gray et al. 2019a) as the computer performed the image-based test considerably faster on average than humans. The image test results are consistent with other deep learning related models comparing human and computer performance. Villon et al. (2018) trained a classification model which outperformed humans by approximately 5% in classifying still images of nine coral reef fish species. Similar results were found by Torney et al. (2019) using object detection to accurately survey wildebeest abundance in Tanzania at a rate of approximately 500 images per hour. Torney et al. (2019) calculated that computer analysis could reduce analysis of surveys from around three to six weeks done manually by up to four wildlife experts, down to just 24 hours using a deep learning algorithm. Additionally, they found accuracy was not compromised, with the abundance estimate from deep learning within 1% of that from expert manual analysis. Like humans, the computer is reliant on the quality of the image it receives. Deep learning methods tend to decrease in performance when the picture quality is blurred or subject to excessive noise (Salman et al. 2016). In low light or high turbidity situations, image processing to improve the quality of the picture (such as cancelling noise and improving contrast) can improve the performance of the model (Salman et al. 2016).

Previous studies comparing humans versus computers have predominantly used images rather than videos. When analysing video footage, there is an assumption that humans have the comparative advantage when addressing uncertainty and ambiguity (Jarrahi 2018). Fish that could not be positively identified early in the video may be identifiable later and vice versa. Humans can move back and forward within the video to correctly identify each fish when calculating MaxN, an ability our deep learning model lacks. The results show that even when humans seem to have the spatio-temporal advantage, the computer model still outperforms both the experts and citizen scientists. In our set-up, inference time for video footage by the computer was about half that of humans. Analytical time could be further reduced by using multiple GPUs or by implementing parallel processing using multiple virtual machines. Consistency in estimating populations is important in ecology, as quantifying population trends is critical to understanding ecosystem health. The computers low variation indicates that it may prove an advantage for monitoring, when data relies on consistency to determine fluctuations in species abundance. Errors that can occur when using humans in data analysis can include individual observer bias and even bias estimation of trends (Yoccoz et al. 2001). This variance is inter-personal and could be standardised by having a single observer across all data sets. This is unrealistic, however, given the large volumes of data often generated by video monitoring (Weinstein 2017). Deep learning methods standardise observer affects not only within data-sets, but also between data-sets from different periods, without personal bias.

The performance curves for our models suggest that they may be just as useful in determining fish abundance with fewer annotations than our full training set of 6,080 annotations. Therefore, less time was needed for training the algorithm as the accuracy of the model’s ability to predict the whole fish (mAP50) is not needed to determine abundance. As our model took approximately 60 hours to train, running a performance curve while training we can see that the time to reach optimum performance could be two-thirds quicker at 20 hours. Creating a performance curve is a useful step when calculating the cost-benefits of implementing a high performing model as well as monitoring algorithm issues such as overfitting. However, this does not take into account the time for human to be trained on which species to annotate. Fish identification experts may not need additional training while citizen scientists may. However studies have shown that citizen scientist annotated data for deep learning can be as reliable as expertly annotated data (Snow et al. 2008) providing an additional low-cost solution for model training.

Although recent advances in deep learning can make image analysis for animal ecology more efficient, there are still some ecological and environmental limitations. Ecological limitations include the difficulty in detection of small, rare or elusive species and therefore abundance may not be able to be estimated in-situ. Nevertheless, even plankton classification using deep learning has been attempted (Li & Cui 2016, Py et al. 2016). This approach may be used to calculate the relative abundance of these microscopic organisms and therefore estimate a wild population density. This may be particularly useful in predicting and monitoring outbreaks of nuisance species such as crown-of-thorns sea stars (Hock et al. 2014) or stinging sea jellies (Llewellyn et al. 2016). Another key ecological issue when using computer vision is low sampling resolution due to the limited field of view from cameras, limiting the accuracy of determining abundance. Campbell et al. (2018) discovered that using cameras with a 360-degree field-of-view improved the accuracy of fish counts compared with single-camera MaxN counts. Improvements for future studies could include combining deep learning with a 360-degree camera aspect when assessing abundance. The current limitations in computer vision imply that this technology is not suitable for all facets of animal ecology. Environmental conditions such as water clarity and light availability currently dictate the useability of footage in marine environments which subsequently affects the performance of the model (Salman et al. 2019). However, these limitations are also experienced by human observers in manual data analysis.

Deep learning methodologies provide a useful tool for consistent monitoring and estimations of abundance in marine environments, surpassing the overall performance of manual, human efforts in a fraction of the time. As this field advances, future ecological applications can include automation in estimating fish size (Costa et al. 2006), estimating abundance for multiple species simultaneously (Mandal et al. 2018), studying animal behaviour (Valletta et al. 2017, Norouzzadeh et al. 2018), and monitoring pest species populations (Clement et al. 2005). Future technological advances in the application of the “internet of things” may also provide ecologists with fully automated management systems via remote sensors connected to machine learning algorithms to achieve continuous environmental information at high temporal resolution (Allan et al. 2018). Given the significant advantages that these algorithms can provide, deep learning can indeed be a highly successful and complementary tool for marine animal ecology.

## Acknowledgements

We thank the many fish experts and citizen scientists who participated in the study. This work benefitted from the support of the Global Wetlands Project.

